# Protease shaving resolves the surface proteome of infection-derived extracellular vesicles and unmasks their inflammatory activity

**DOI:** 10.64898/2026.11.22.768461

**Authors:** Jorge J. Canas, Claire Laurila, Samantha M. Enslow, Xin Wang, Mariola J. Ferraro

## Abstract

Small extracellular vesicles (sEVs) released by infected macrophages carry both host proteins and bacterial antigens, but whether these components face outward, are enclosed within vesicles, or are associated with co-isolated material remains unclear, limiting our understanding of their accessibility to recipient cells. We combined proteinase K shaving with quantitative proteomics to characterize the protease-accessible proteome of sEVs released by *Salmonella*-infected RAW264.7 macrophages. Host and bacterial proteins exhibited distinct responses to proteolysis. Of 1,368 proteins quantified, 246 were significantly decreased following digestion and a further 93 became undetectable. Among the depleted proteins, 26 carried independent topological evidence of surface exposure, spanning tetraspanins, integrins, MHC class I molecules, GPI-anchored proteins and membrane receptors, and the depleted fraction also included the immune mediators TNF-α, CSF2RA, CD40 and MIF. Most detected *Salmonella* proteins resisted proteolysis and seven were significantly increased. Peptide-level analysis localized TNF-α depletion to its extracellular domain, which fell by more than 3 log₂ relative to a cytoplasmic peptide measured in the same injection. Preservation of luminal sEV markers and two normalization-independent measures indicated that vesicle integrity was maintained. Despite removal of this protease-accessible fraction, vesicle bioactivity was not diminished: proteinase K-treated vesicles induced stronger *Il6* expression and NF-κB/AP-1 signaling than untreated vesicles, with both responses sensitive to TNF-α blockade and polymyxin B. Our findings together suggest that the protease-accessible protein fraction influences the immunostimulatory activity of infection-derived sEV preparations, with removal of the externally available coat of EVs being associated with enhanced inflammatory signaling.

## Introduction

Cells secrete extracellular vesicles (EVs) during homeostasis and infection to facilitate communication between donor and recipient cells. EVs are lipid bilayer-enclosed particles that carry bioactive molecules associated with their membranes or enclosed within their lumen, including nucleic acids, lipids, metabolites, and proteins ^1^. These EV cargos can influence immunological processes, including cellular targeting, coordination of host defense, and antigen transfer ^2,3^. The cellular origin of EVs can also determine their molecular composition and their interactions with specific recipient cell populations ^1^. In the broad category of EVs, small extracellular vesicles (sEVs) carry a broad range of receptors, ligands, cytokines, and antigens ^4–8^, which can include pathogen-encoded proteins ^6^. Proteins associated with sEV preparations may be exposed on the vesicle surface, present on a corona ^9^, incorporated into the vesicle membrane, enclosed within a protease-protected lumen, and finally associated with non-vesicular proteinaceous material^10,11^, which inadvertently may be present in the sEV preparations despite multiple purification steps being performed ^4^. This topology is likely to influence protein accessibility to cellular receptors and other molecules, and, consequently, how individual cargo molecules interact with recipient cells.

Macrophages are central mediators of innate immune defense, inflammatory responses, and the signaling events that shape subsequent adaptive immunity ^12,13^. During infection, macrophages release sEVs containing host-derived molecules, including cytokines, as well as components originating from invading microorganisms. Tumor necrosis factor alpha (TNF-α) is a pleiotropic cytokine with major roles in inflammation, immune regulation, and cell death ^14–16^. TNF-α typically exists as a 26-kDa membrane-associated precursor that can be proteolytically processed by proteases to generate a 17-kDa soluble form ^17,18^. During bacterial infection, while the host and pathogen interact, macrophages can package bacterial components along with host-derived molecules into sEVs, providing a potential mechanism for disseminating microbial antigens and inflammatory signals beyond the infected cell, potential decoy mechanisms and even potential pathogenic mechanism ^5,19–24^.

Gram-negative intracellular pathogens like *Salmonella enterica* express virulence factors that alter host immune responses to promote intracellular survival, persistence, and dissemination^25^. Previous work has shown that *Salmonella*-derived proteins and pro-inflammatory and other host factors are associated with sEVs released from infected macrophages^6,26–28^ and that *Salmonella* proteins can be also carried by sEVs from other cell types ^29^. How these various components are arranged within sEV preparations, however, remains unknown. Specifically, it is unclear which host and bacterial proteins are displayed on the vesicle surface, and which are shielded within a protected compartment. The distinction matters because surface-accessible and protected proteins differ in their availability to the receptors, recipient-cell lectins, external proteases and antibodies that vesicles may encounter after release.

In this study, we investigated how host proteins and bacterial antigens are distributed between the surface-accessible and protease-protected compartments of sEV preparations from *Salmonella*-infected macrophages at 24 hours post-infection in vitro. We compared commonly used approaches for disrupting or removing surface-associated proteins and identified proteinase K digestion as the most effective in this setting for removing surface-accessible epitopes while preserving vesicle size and concentration. Applying this approach with quantitative proteomics, we asked three questions: which host proteins are externally accessible, whether bacterial antigens share that accessibility, and whether removing the accessible protein layer alters the immunostimulatory activity of the sEVs. Since proteolysis removes a large fraction of total protein and complicates relative quantification, we also used normalization-independent measures to assess vesicle integrity. We found that proteinase K depleted a substantial fraction of host proteins, including established EV surface proteins and immune-related mediators, while most detected *Salmonella* proteins remained resistant to digestion, suggesting they remain in the lumen of sEVs. Peptide-level analysis localized the loss of vesicle-associated TNF-α to its extracellular domain, while removal of the protease-accessible protein fraction enhanced rather than diminished inflammatory signaling in recipient macrophages.

## Results

### Proteinase K outperforms physical disruption for removing surface epitopes

Small extracellular vesicles, sEVs, were isolated from *Salmonella*-infected and uninfected RAW264.7 macrophages (MOI 5:1, 24 hpi, hours post-infection) by differential centrifugation, 0.22 µm filtration and ultracentrifugation, then characterized by nanoparticle tracking analysis (**Fig. 1A)**. To select a method for exposing surface-associated epitopes, we compared probe sonication, heat denaturation and proteinase K digestion across two independent preparations from each condition (infected or uninfected macrophage cultures), profiling equal numbers of vesicles by multiplex bead-based flow cytometry. Among the tested methods, proteinase K was consistently the most disruptive, reducing CD63, CD81 and CD40 significantly more than either physical method in uninfected vesicles, and CD63 and CD81 in infected vesicles, while sonication and heat denaturation were effective, but they were indistinguishable from one another for tested markers (**Fig. S1A**). Nanoparticle tracking analysis confirmed that proteinase K did not significantly alter the size or concentration of treated sEVs (**Fig. S1B, C**), indicating that the reduction in marker signal reflected enzymatic removal of accessible proteins rather than disruption or loss of vesicles. Proteinase K treatment is also well established for defining EV surface proteins ^30,31^.

**Figure 1.**
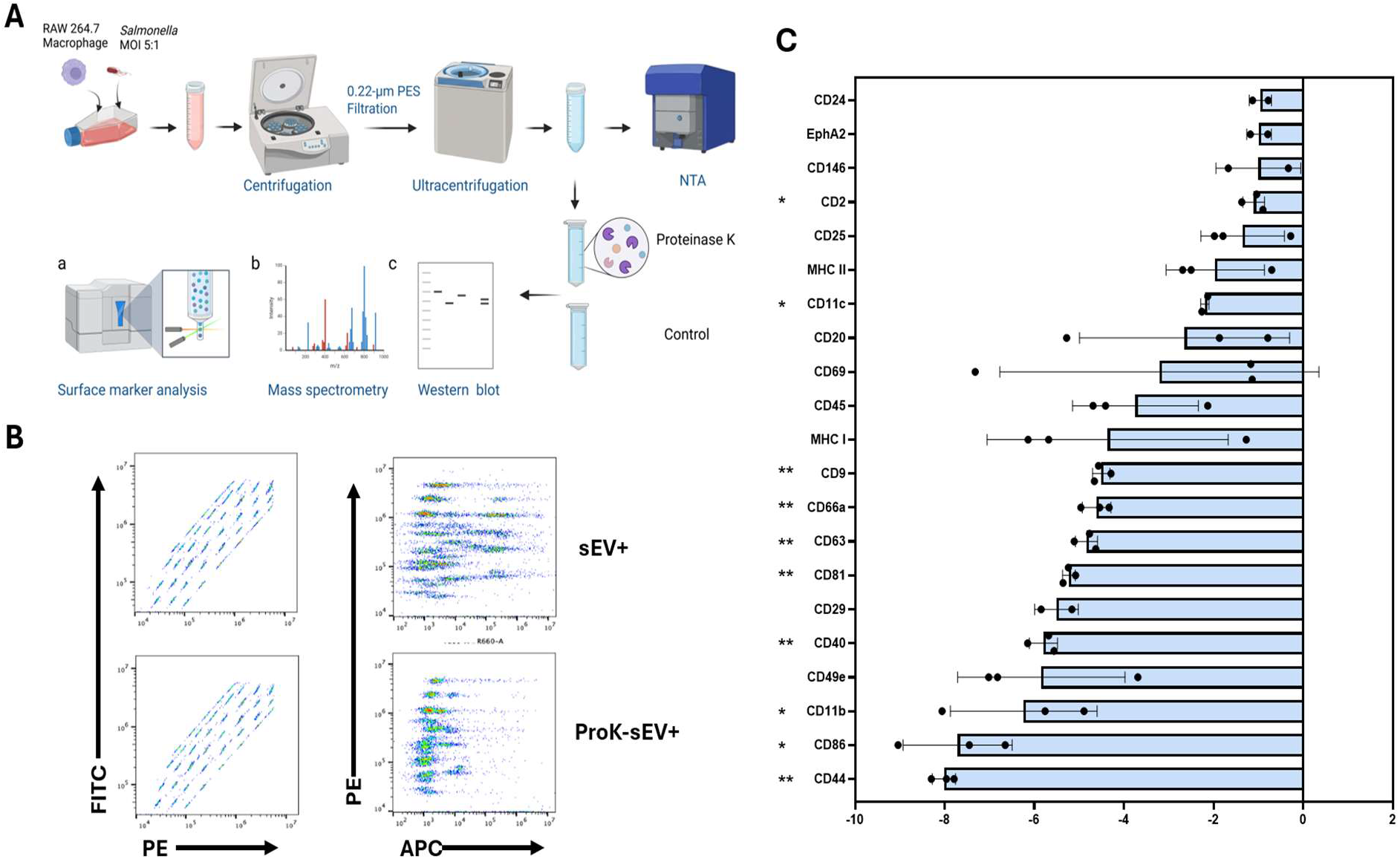
Proteinase K shaving of sEVs removes surface-accessible epitopes from vesicles released by *Salmonella*-infected macrophages. (A) Experimental workflow. sEVs were isolated from RAW264.7 macrophages infected with *Salmonella* Typhimurium (MOI 5:1, 24 hpi) or left uninfected, by differential centrifugation, 0.22-µm filtration and ultracentrifugation; characterized by nanoparticle tracking analysis (NTA); and treated with proteinase K or buffer alone before analysis by multiplex bead-based surface profiling, mass spectrometry and Western blotting. (B) Representative flow cytometry of the multiplex bead panel for untreated and proteinase K-treated sEV(+), showing bead population resolution (FITC versus PE) and marker detection (PE versus APC). (C) Log₂ fold change in APC signal after proteinase K treatment relative to untreated sEV(+) (n = 3 independent digestions from two biological preparations). Markers whose untreated signal fell below twice the buffer control were excluded. Bars show mean ± SD with individual digestions overlaid. Significance was assessed by two-tailed one-sample *t*-test against a null log₂ fold change of zero, with Benjamini–Hochberg correction; q < 0.05 (*), q < 0.01 (**).

### Proteinase K broadly reduces surface-accessible epitopes on infected-cell sEVs

Because our main objective was to characterize sEVs released during bacterial infection, subsequent experiments focused on vesicles from *Salmonella*-infected RAW264.7 macrophages only [sEV(+); MOI 5:1, 24 hpi; **Fig. 1A**], analyzed by multiplex bead-based surface profiling, mass spectrometry and Western blotting. We used multiplex bead-based flow cytometry to determine which host surface markers were detectable on sEVs from infected macrophages and to assess their accessibility to proteinase K. Bead populations were clearly resolved in the FITC and PE channels and remained unaffected by digestion, indicating that proteinase K treatment did not interfere with bead identification. In contrast, the APC detection signal was markedly reduced across most bead populations following treatment (**Fig. 1B**). Detection thresholds were defined individually for each bead population as twice the signal of the corresponding buffer-only control. Eight bead populations, CD3, CD4, CD49b, CD105, AN2, CD41, CD61 and the rat IgG1 isotype control, did not exceed this threshold even in untreated sEV preparations and were therefore considered below the limit of quantification. The absence of detectable signal for the isotype control supported the specificity of the assay threshold.

Among markers detectable on untreated sEV(+), proteinase K treatment substantially reduced APC signals across the panel (**Fig. 1C**). Ten markers remained significantly reduced after Benjamini–Hochberg correction: CD44 (log₂FC ≤ −8.02; ≤0.4% of untreated; q = 0.004), CD86 (−7.71; 0.5%; q = 0.026), CD11b (−6.23; 1.3%; q = 0.049), CD40 (−5.79; 1.8%; q = 0.004), CD81 (≤−5.23; ≤2.7%; q = 0.004), CD63 (−4.83; 3.5%; q = 0.004), CD66a (≤−4.61; ≤4.1%; q = 0.006), CD9 (−4.51; 4.4%; q = 0.004), CD11c (≤−2.20; ≤21.8%; q = 0.048), and CD2 (−1.10; 46.5%; q = 0.043). Notably, all three canonical EV-associated tetraspanins, CD9, CD63, CD81, showed reductions to less than 5% of their untreated signals, with CD81 falling below the detection threshold in every proteinase K-treated sample.

### Proteomic analysis identifies a broad surface-accessible host protein fraction

To extend our analysis beyond the predefined marker panel, we characterized the protease-accessible proteome of sEV(+) using quantitative mass spectrometry, comparing proteinase K-treated vesicles with untreated controls. Following filtering, 9,732 peptide groups mapping to 2,076 protein groups were quantified, of which 1,368 were supported by minimum two peptides to avoid one-hit wonders, and retained for analysis. Proteinase K treatment resulted in 246 significantly decreased and 197 significantly increased proteins (Benjamini–Hochberg-adjusted *P* < 0.05; absolute fold change ≥ 1.5). An additional 93 proteins were assigned to a separate “no longer detected after ProtK” category based on exclusively missing-not-at-random-classified peptides and negative estimated log₂ fold changes. Because these estimates depended on detection-limit imputation, these proteins were reported separately from the 246 significantly decreased proteins. Combining the significantly decreased and no-longer-detected categories identified 339 proteins with reduced detectability following proteinase K treatment (**Fig. 2A; Tables S1–S3**).

**Figure 2.**
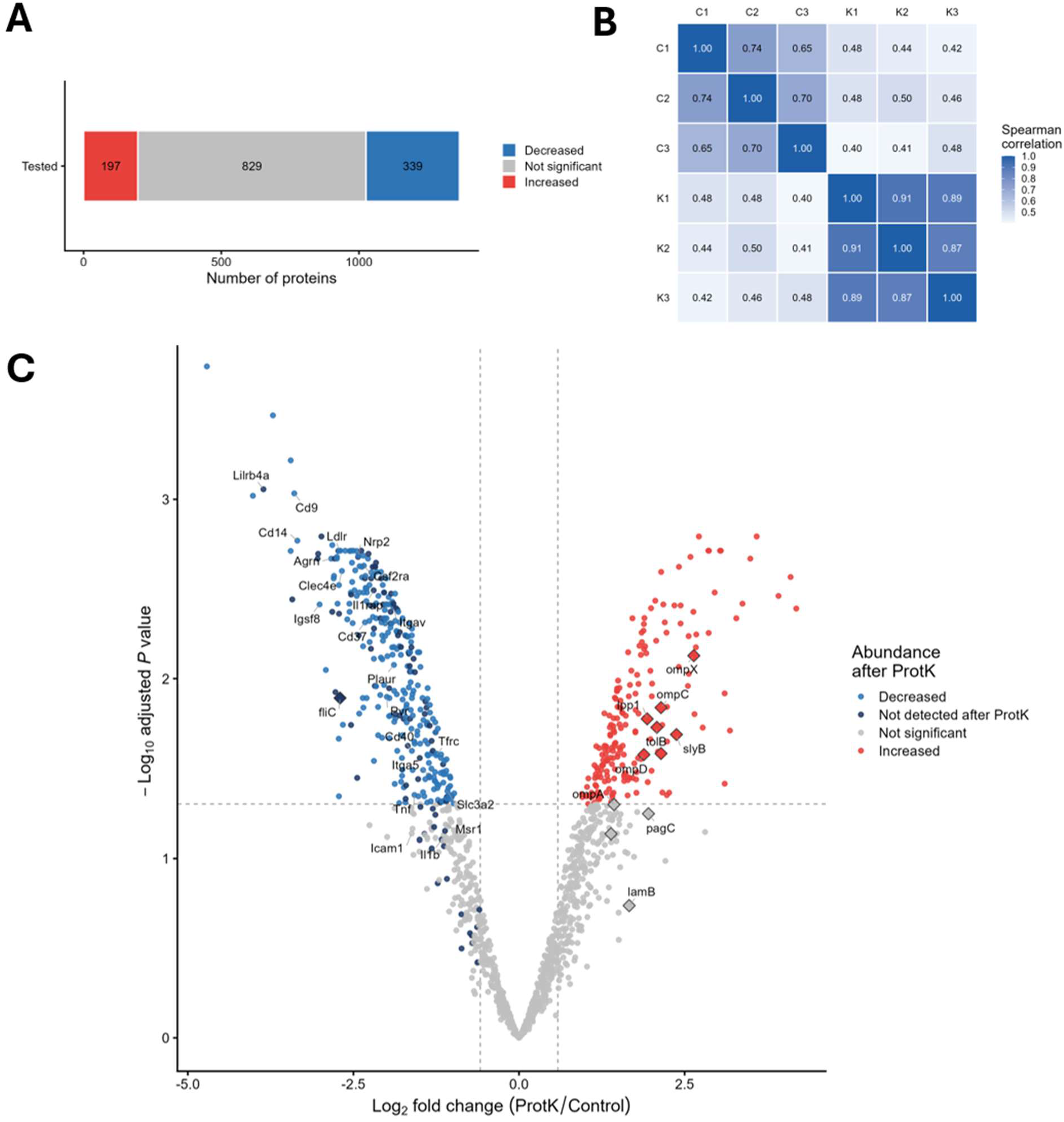
Proteinase K shaving identifies the surface-accessible proteome of sEVs from *Salmonella*-infected cells. sEVs from infected RAW264.7 macrophages [sEV(+)] were left untreated (Control) or treated with proteinase K (ProtK) before LC–MS/MS analysis (n = 3 per condition). (**A**) Differential abundance outcome for the 1,368 proteins supported by at least two peptides: decreased (n = 339, comprising 246 significantly decreased and 93 detected in no treated replicate), not significant (n = 829) and increased (n = 197). Three proteins detected in no control replicate were excluded as imputation artefacts, so the categories sum to 1,365. (**B**) Spearman correlation between all pairs of samples, calculated on median-centred log₂ protein abundances (**C**) Volcano plot showing log₂ fold change against −log₁₀ adjusted *P*. Dashed lines indicate a 1.5-fold change and adjusted *P* = 0.05. Circles, mouse proteins; diamonds, *Salmonella* Typhimurium proteins; dark blue, proteins detected in no treated replicate. Selected proteins are labelled.

Proteinase K treatment was the major source of variation in the proteomic dataset. Treated replicates showed strong within-group correlations (Spearman’s ρ = 0.87–0.91), whereas correlations between treated and untreated samples were substantially lower (ρ = 0.40–0.50; **Fig. 2B**).

Proteins depleted following proteinase K treatment included numerous established EV surface-associated proteins. These included the tetraspanins Cd9 (log₂FC = −3.39), Igsf8 (−3.01), and Cd37 (−2.33); the GPI-anchored proteins Cd14 (−3.35) and Plaur (−1.89); the receptors Lilrb4a (−3.86), Ldlr (−2.72), Nrp2 (−2.43), Csf2ra (−2.34), Cd40 (−1.62), and Tfrc (−1.25); and the integrin subunits Itgav (−1.90) and Itga5 (−1.36). (**Fig. 2C**, **Table 1, Table S1**).

**Table 1.**
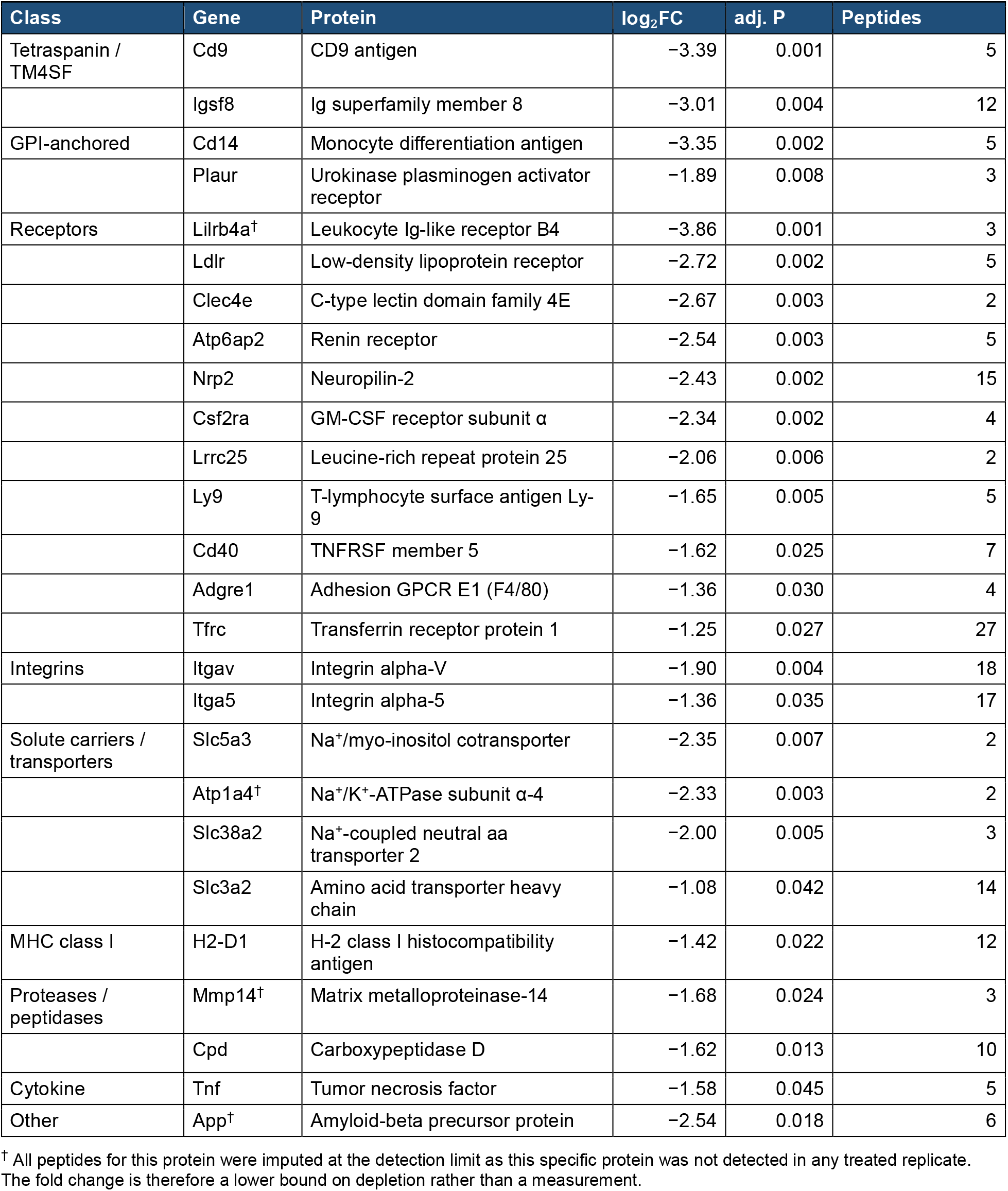
Surface-exposed proteins depleted by proteinase K treatment of sEV(+). All proteins with reduced abundance or detectability after treatment that carry UniProt topological evidence of surface exposure: 22 significantly decreased (adj. *P* < 0.05, fold change ≥ 1.5, ≥2 peptides) and four no longer detected after digestion. Proteins are grouped by functional class and ordered by magnitude within each group.

| Class | Gene | Protein | log<sub>2</sub>FC | adj. P | Peptides |
|---|---|---|---|---|---|
| Tetraspanin / TM4SF | Cd9 | CD9 antigen | −3.39 | 0.001 | 5 |
|  | Igsf8 | Ig superfamily member 8 | −3.01 | 0.004 | 12 |
| GPI-anchored | Cd14 | Monocyte differentiation antigen | −3.35 | 0.002 | 5 |
|  | Plaur | Urokinase plasminogen activator receptor | −1.89 | 0.008 | 3 |
| Receptors | Lilrb4a<sup>†</sup> | Leukocyte Ig-like receptor B4 | −3.86 | 0.001 | 3 |
|  | Ldlr | Low-density lipoprotein receptor | −2.72 | 0.002 | 5 |
|  | Clec4e | C-type lectin domain family 4E | −2.67 | 0.003 | 2 |
|  | Atp6ap2 | Renin receptor | −2.54 | 0.003 | 5 |
|  | Nrp2 | Neuropilin-2 | −2.43 | 0.002 | 15 |
|  | Csf2ra | GM-CSF receptor subunit α | −2.34 | 0.002 | 4 |
|  | Lrrc25 | Leucine-rich repeat protein 25 | −2.06 | 0.006 | 2 |
|  | Ly9 | T-lymphocyte surface antigen Ly-9 | −1.65 | 0.005 | 5 |
|  | Cd40 | TNFRSF member 5 | −1.62 | 0.025 | 7 |
|  | Adgre1 | Adhesion GPCR E1 (F4/80) | −1.36 | 0.030 | 4 |
|  | Tfrc | Transferrin receptor protein 1 | −1.25 | 0.027 | 27 |
| Integrins | Itgav | Integrin alpha-V | −1.90 | 0.004 | 18 |
|  | Itga5 | Integrin alpha-5 | −1.36 | 0.035 | 17 |
| Solute carriers / transporters | Slc5a3 | Na<sup>+</sup>/myo-inositol cotransporter | −2.35 | 0.007 | 2 |
|  | Atp1a4<sup>†</sup> | Na<sup>+</sup>/K<sup>+</sup>-ATPase subunit α-4 | −2.33 | 0.003 | 2 |
|  | Slc38a2 | Na<sup>+</sup>-coupled neutral aa transporter 2 | −2.00 | 0.005 | 3 |
|  | Slc3a2 | Amino acid transporter heavy chain | −1.08 | 0.042 | 14 |
| MHC class I | H2-D1 | H-2 class I histocompatibility antigen | −1.42 | 0.022 | 12 |
| Proteases / peptidases | Mmp14<sup>†</sup> | Matrix metalloproteinase-14 | −1.68 | 0.024 | 3 |
|  | Cpd | Carboxypeptidase D | −1.62 | 0.013 | 10 |
| Cytokine | Tnf | Tumor necrosis factor | −1.58 | 0.045 | 5 |
| Other | App<sup>†</sup> | Amyloid-beta precursor protein | −2.54 | 0.018 | 6 |

<sup>†</sup> All peptides for this protein were imputed at the detection limit as this specific protein was not detected in any treated replicate. The fold change is therefore a lower bound on depletion rather than a measurement.

To assess whether protein depletion was consistent with surface exposure, we examined the membrane topology of the depleted proteins. Twenty-six proteins had supporting topological annotations in UniProt, including a GPI anchor or a transmembrane segment with an annotated extracellular domain (**Fig. 3A**). These represented several classes of membrane-associated proteins, such as tetraspanins Cd9 and Igsf8; the GPI-anchored proteins Cd14 and Plaur; the receptors Lilrb4a, Ldlr, Nrp2, Csf2ra, Cd40, Ly9, Tfrc, and Clec4e; the integrin subunit Itgav; the solute carriers Slc5a3, Slc38a2, and Slc3a2; the macrophage marker Adgre1 (F4/80); MHC class I protein H2-D1; and TNF-α. Their log₂ fold changes ranged from −1.08 for Slc3a2 to −3.86 for Lilrb4a, the latter a bound since Lilrb4a was not detected after treatment, with 2–27 peptides identified per protein. Cd37 showed a comparable decrease (log₂FC = −2.33) but was not included among the 26 proteins because its UniProt entry lacked the extracellular topology annotation required for inclusion (**Fig. 3B)**.

**Figure 3.**
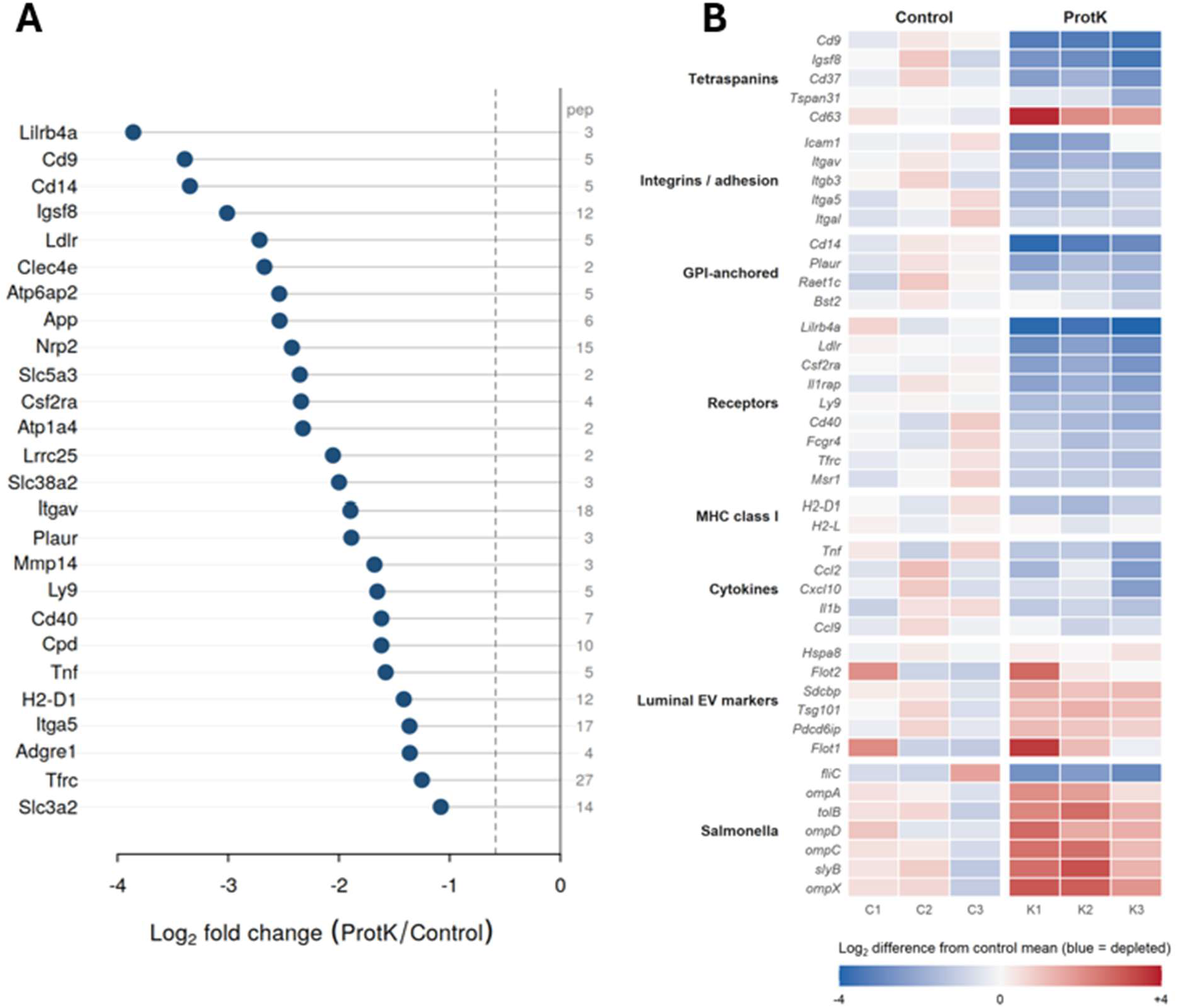
Surface-annotated proteins are selectively depleted while luminal and bacterial proteins are retained. (**A**) Twenty-six proteins showing reduced abundance or detectability after proteinase K treatment and carrying UniProt topological evidence of surface exposure: 22 significantly decreased proteins and four classified as no longer detected. Proteins are ranked by estimated log₂ fold change; these estimates represent detection-limit-dependent bounds for proteins in the latter category. Peptide counts are shown at right. (**B**) Log₂ abundance difference from the mean of untreated controls for selected proteins, grouped by functional class, each replicate shown.

#### Luminal proteins and bacterial outer membrane proteins were retained in EVs post-proteinase K digestion

Because proteinase K removes a substantial fraction of total protein, relative abundance after normalization cannot by itself establish that vesicles remained “sealed”. We therefore applied two normalization-independent measures. First, luminal EV proteins maintained similar positions within the sample abundance distributions before and after digestion (86.5th–87.9th versus 87.9th–90.2nd percentile), while surface-associated proteins shifted markedly downward (65.8th–74.3rd to 38.2nd–44.9th percentile; **Fig. S2A**). Second, 85.2% of luminal peptides detected in at least two untreated replicates remained detectable after digestion, compared with 51.6% of surface-associated peptides. Retention was high for Pdcd6ip (92%), Rab7a (91%), Tsg101 (89%), and Hspa8 (84%), but substantially lower for Lilrb4a (0%), Il1rap (0%), Agrn (7%), and Cd37 (20%; **Fig. S2B**). These findings support preferential preservation of luminal proteins and are consistent with substantial maintenance of vesicle integrity during proteolysis.

Most detected *Salmonella* proteins were retained following digestion, and seven were significantly increased relative to the surviving proteome: OmpX (log₂FC = +2.64), SlyB (+2.38), OmpC (+2.14), YncE (+2.14), TolB (+2.08), Lpp1 (+1.94) and OmpD (+1.89), all adjusted *P* < 0.05. Their resistance to proteolysis is consistent with protection within vesicles or intrinsic protease resistance, while the increases are most plausibly compositional, reflecting a larger share of the protein remaining after accessible host protein was removed.

Since proteomic results often depend on the normalization scheme, we compared eight normalization schemes to assess the sensitivity of differential abundance results to sample-level scaling (**Table S5, Fig. S3**). Protein rankings were identical across schemes (Spearman’s ρ = 1.000), as each applied a sample-level offset. Under these alternative global scaling methods, 237–246 of the 246 proteins classified as decreased using median centering retained this classification. In contrast, normalization to a luminal EV reference shifted log₂ fold changes by −0.99 relative to median centering, increasing the number of proteins classified as decreased from 246 to 488 and reducing those classified as increased from 197 to 28. The 93 proteins classified as no longer detected remained still unchanged. Therefore, although protein rankings were preserved, estimated fold changes and differential abundance classifications depended on the normalization reference. However, median centering was retained as the primary normalization method, with alternative schemes used to assess sensitivity.

#### Vesicle-associated TNF-α exhibits proteolytic accessibility consistent with membrane-anchored topology

TNF-α was consistently detected in untreated sEV(+) preparations and significantly depleted following proteinase K treatment (log₂FC = −1.58, adj. *P* = 0.045; **Figs. 2C and 3A**), indicating substantial protease accessibility. To determine if this susceptibility followed the expected topology of membrane-anchored murine pro-TNF-α, we examined five unique peptides mapping to TNF-α’s cytoplasmic and extracellular domains (**Fig. 4A; Table S4**).

**Figure 4.**
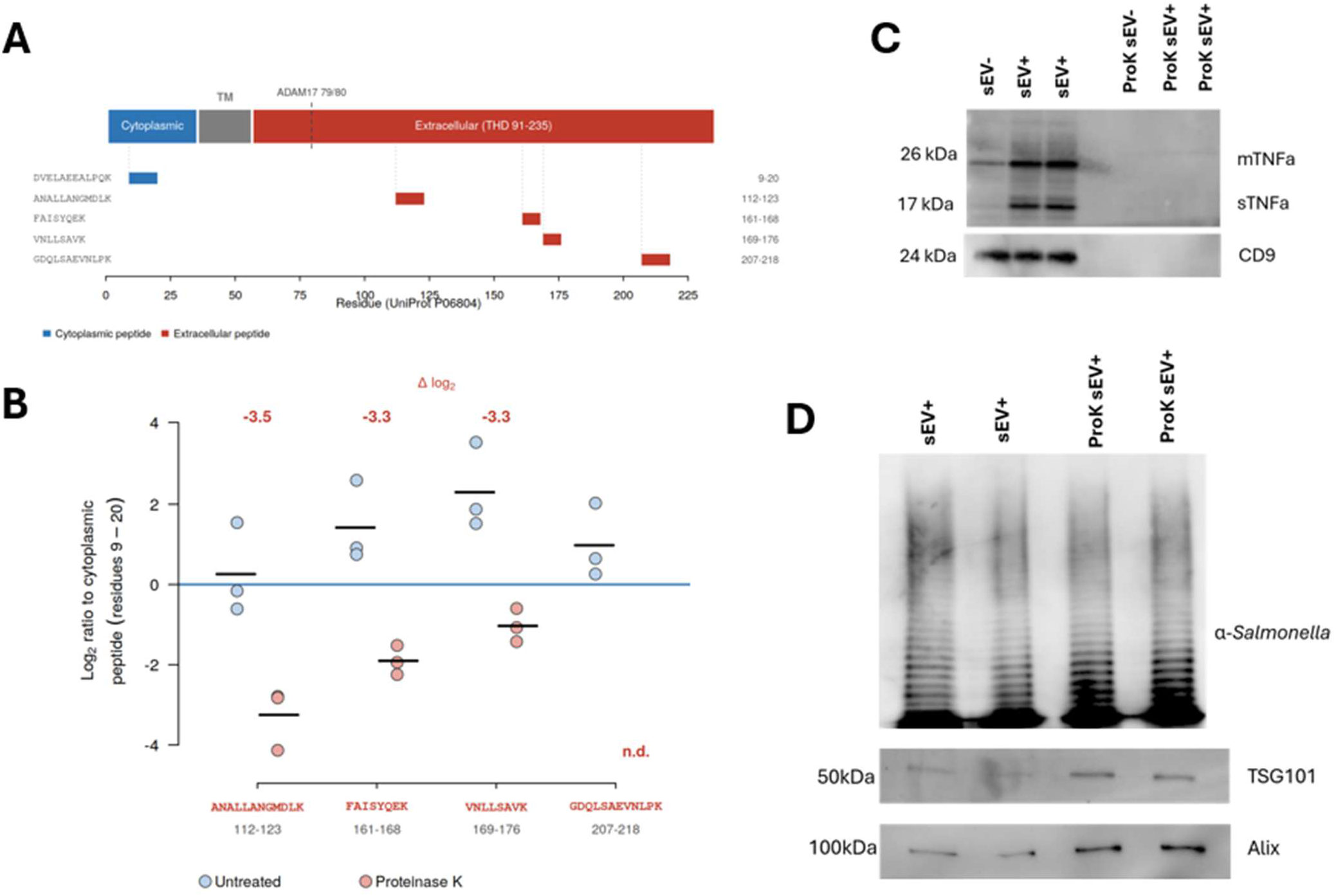
TNF-α is protease accessible whereas *Salmonella* antigens and luminal EV proteins are protected from the proteinase K treatment. (**A**) Topology of mouse pro-TNF-α (UniProt P06804): cytoplasmic domain (residues 1–35), transmembrane segment (36–56) and extracellular domain (57–235), with the positions of the five quantified tryptic peptides and the ADAM17 cleavage site. (**B**) Each extracellular peptide expressed as a log₂ ratio to the cytoplasmic peptide DVELAEEALPQK in the same injection (n = 3; bars, mean), with the mean difference between conditions above each peptide. n.d., not detected. (**C**) Untreated and proteinase K-treated (ProtK) sEVs from uninfected [sEV(−)] and infected [sEV(+)] macrophages, probed for TNF-α and CD9. mTNFα, 26-kDa membrane-associated form; sTNFα, 17-kDa soluble form. (**D**) Untreated and ProtK-treated sEV(+), probed for *Salmonella* O/H antigen and the luminal markers TSG101 and ALIX. sEV lanes in (C) and (D) were loaded by particle number, 4.5 × 10⁹ vesicles per lane.

The cytoplasmic peptide (residues 9–20) of TNF-α was identified by MS/MS in all three proteinase K-treated replicates and was essentially unchanged in abundance. In contrast, extracellular peptides were strongly depleted: residues 161–168 were identified by MS/MS in two treated replicates, residues 112–123 and 169–176 were quantified by precursor intensity without MS/MS confirmation in the treated samples, and residues 207–218 became undetectable. Relative to the cytoplasmic peptide measured within the same injection, the three remaining quantifiable extracellular peptides decreased by 3.3–3.5 log₂ units (**Fig. 4B**). These normalization-independent ratios show preferential depletion of the extracellular TNF homology domain relative to the N-terminal cytoplasmic region. This pattern is consistent with the type II transmembrane topology of murine pro-TNF-α, in which the extracellular domain is exposed to proteolysis while the cytoplasmic region is relatively protected by the membrane.

Immunoblotting further demonstrated loss of detectable full-length (26 kDa) and soluble (17 kDa) TNF-α following digestion, together with CD9 (**Fig. 4C**). Since the antibody used in this study recognizes epitopes within the mature extracellular TNF-α domain, loss of immunoreactivity is consistent with cleavage of this region and does not exclude persistence of an N-terminal membrane-associated fragment. The 17-kDa form was detected only in infected sEV preparations and was also present in infected-cell pellets and conditioned medium (**Fig. S4A**). Its presence is consistent with ectodomain shedding of membrane-associated TNF-α, a process primarily mediated by ADAM17, which was itself significantly depleted by proteinase K in our proteome (log₂FC = −1.13, adj. *P* = 0.029; **Table S1**), indicating that both the sheddase and its substrate are externally accessible.

Although the presence and protease sensitivity of soluble TNF-α do not establish its vesicle-surface localization, the preferential retention of the cytoplasmic peptide alongside pronounced extracellular-domain depletion supports a membrane-anchored orientation of pro-TNF-α in infected sEV preparations (**Fig. S4B**), with an externally accessible extracellular domain and a relatively protected N-terminal region.

### Differential protease sensitivity of host immune mediators and bacterial antigens

Protease accessibility extended to several additional immune-related proteins, but responses varied across the proteins examined (**Table 2**). The receptor subunits Csf2ra (log₂FC = −2.34, adjusted *P* = 0.002) and Cd40 (−1.62, adj. *P* = 0.025), as well as Mif (−1.24, adjusted *P* = 0.026), were significantly depleted following digestion. Il1rap and Il1b were detected in all three untreated replicates but became undetectable following proteinase K treatment and were thus reported as no longer detected rather than assigned numerical fold changes. In contrast, Ccl2 (−1.50), Cxcl10 (−1.21), and Ccl9 (−0.55) showed decreases that did not reach statistical significance, while Ccl5 showed no significant change (+0.56). These findings indicate that protease sensitivity varied among immune-related proteins. Ebi3, Sdc4, Cd276, and Il1rn (IL-1Ra) were each supported by only one peptide and therefore excluded from the reported protein set, with their apparent depletion considered provisional.

**Table 2.** Cytokines, chemokines and their receptors detected in sEV(+) preparations. All detected immune mediators and receptor subunits supported by at least two peptides, ordered by log_2_ fold change.

| Gene | Protein | log<sub>2</sub>FC | adj. P | Peptides | Classification |
|---|---|---|---|---|---|
| Csf2ra | GM-CSF receptor subunit α | −2.34 | 0.002 | 4 | Decreased |
| Il1rap<sup>†</sup> | IL-1 receptor accessory protein | −2.20 | 0.003 | 3 | Not detected after ProtK |
| Cd40 | TNFRSF member 5 | −1.62 | 0.025 | 7 | Decreased |
| Tnf | Tumor necrosis factor | −1.58 | 0.045 | 5 | Decreased |
| Ccl2 | C-C motif chemokine 2 | −1.50 | 0.092 | 2 | Not significant |
| Mif | Macrophage migration inhibitory factor | −1.24 | 0.026 | 3 | Decreased |
| Cxcl10 | C-X-C motif chemokine 10 | −1.21 | 0.132 | 3 | Not significant |
| Il1b<sup>†</sup> | IL-1β | −1.17 | 0.079 | 2 | Not detected after ProtK |
| Ccl9 | C-C motif chemokine 9 | −0.55 | 0.299 | 2 | Not significant |
| Csf1r | M-CSF 1 receptor | −0.29 | 0.625 | 7 | Not significant |
| Tnfrsf1b | TNFRSF member 1B | +0.11 | 0.877 | 8 | Not significant |
| Ccl5 | C-C motif chemokine 5 | +0.56 | 0.749 | 7 | Not significant |

<sup>†</sup> Detected in all three untreated replicates and in none of the treated replicates. All peptides were imputed at the detection limit, and so the fold change is a lower bound on depletion.

Unlike several host immune-related proteins, the *Salmonella* antigens examined remained detectable following proteinase K treatment, as did the luminal EV-associated proteins Tsg101 and Pdcd6ip (ALIX; **Fig. 4D**). Because equal particle numbers rather than equal protein amounts were loaded for immunoblotting, preservation of these luminal markers supports retention of vesicle-associated internal proteins despite extensive proteolysis. Together with the marked loss of detectable CD9 and TNF-α, these findings are consistent with preferential digestion of externally accessible proteins while substantial vesicle integrity was maintained. Consistent with the proteomic analysis (**Figs. 2C, 3B**), the bacterial antigens examined were largely resistant to digestion, potentially reflecting protection within membrane-bound compartments or intrinsic resistance to proteolysis.

### Removal of surface-accessible proteins enhances sEV-induced inflammatory signaling

Having established that proteinase K substantially depleted surface-associated host proteins from sEV(+) preparations, we next investigated whether these proteins contribute to or limit the immunostimulatory activity of the vesicles. Untreated and proteinase K-treated sEV(+) preparations were administered at equal protein mass (1 µg per treatment), with protein content re-quantified following digestion and filtration to remove liberated peptide fragments. Following 4 h of stimulation, naïve macrophages exposed to proteinase K-treated sEV(+) showed approximately 4.5-fold greater *Il6* expression than those receiving untreated sEVs(+) (**Fig. 5A**). NF-κB/AP-1 reporter activity measured after 24 h was similarly enhanced (**Fig. 5B**). Thus, removal of the protease-accessible protein fraction was associated with increased rather than diminished inflammatory activity of sEV(+). The flow-through collected following digestion showed no detectable stimulatory activity above background (**Fig. 5B**), indicating that the enhanced response was associated with the retained vesicle-enriched fraction rather than the released digestion products. Because TNF-α was among the protease-sensitive components of sEV(+), we investigated its contribution to the inflammatory response. TNF-α neutralization reduced *Il6* induction by proteinase K-treated sEV(+) to levels comparable to those induced by untreated vesicles, while having no significant effect on the response to untreated sEVs(+) (**Fig. 5A**). We also examined the contribution of bacterial lipid-associated components using polymyxin B, which binds the lipid A moiety of lipopolysaccharide (LPS). Polymyxin B abolished the enhanced *Il6* response induced by proteinase K-treated sEVs(+) (**Fig. 5C**). The response to untreated sEV(+) was also slightly reduced, although this difference did not reach statistical significance. Together, these findings show that proteinase K treatment enhances the inflammatory activity of sEV(+) preparations, with the increased response sensitive to both TNF-α neutralization and polymyxin B treatments.

**Figure 5.**
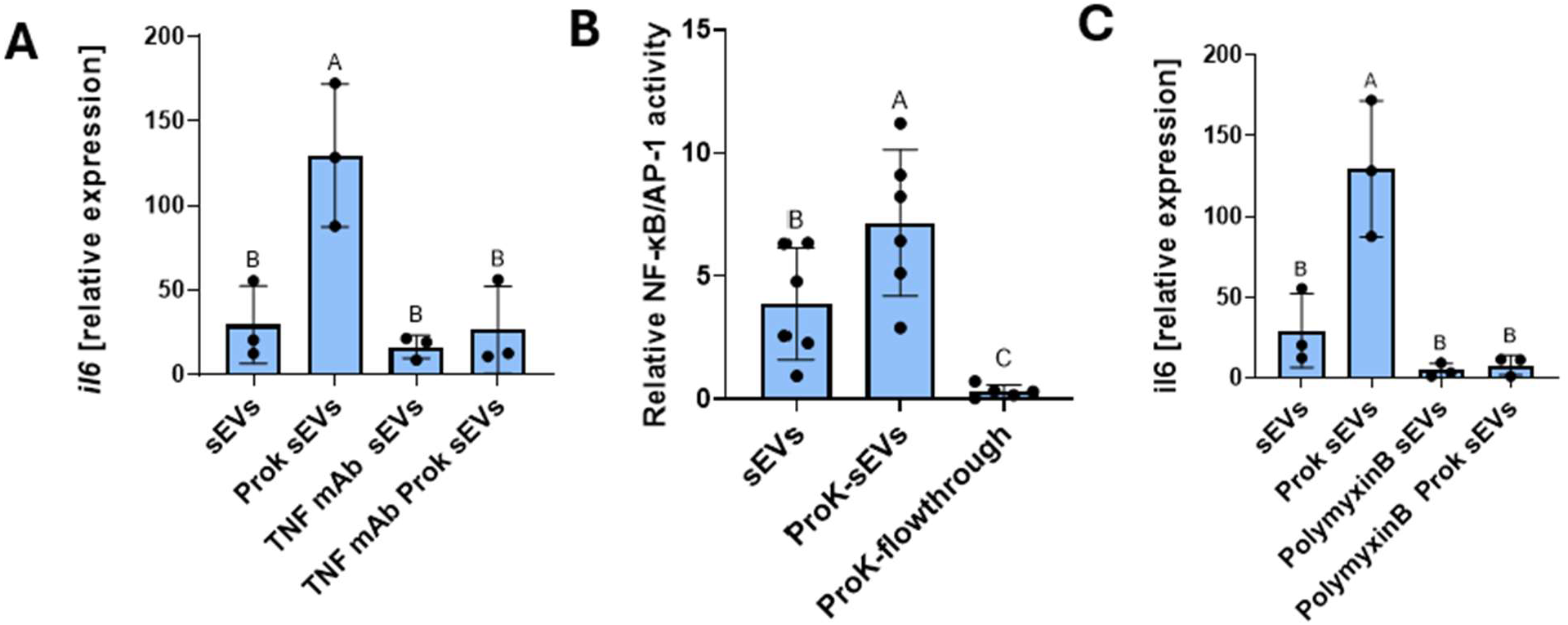
Removal of surface-accessible proteins enhances sEV-induced inflammatory signaling. **(A)** *Il6* expression in naïve RAW264.7 macrophages 4 h after stimulation with 1 µg untreated or proteinase K-treated (ProtK) sEVs(+), with or without TNF-neutralizing antibody. **(B)** NF-κB/AP-1 reporter activity 24 h after stimulation with untreated sEVs(+), ProtK-treated sEVs(+), or the flow-through recovered after digestion. **(C)** *Il6* expression as in (A), with or without polymyxin B. The untreated and ProtK-treated groups are shared between (A) and (C). Bars show mean ± SD with individual replicates overlaid (n = 3 in A and C, n = 6 in B). Groups not sharing a letter differ significantly (one-way ANOVA with Tukey’s post hoc test were applied, p < 0.05).

## Discussion

Small extracellular vesicles (sEVs) have emerged as important mediators of intercellular communication, facilitating the transfer of proteins, cytokines, and damage- or pathogen-associated molecular patterns between cells ^5,32^. Beyond the molecular composition of these vesicles, the spatial distribution and accessibility of their associated components are likely to influence how they interact with recipient cells and regulate immune responses. Proteins exposed on the vesicle exterior may participate directly in receptor engagement and signaling, whereas luminal cargo generally requires vesicle internalization or membrane fusion to access intracellular pathways. Distinguishing these compartments can help us understand the functional properties of sEVs. Consistent with the MISEV guidelines, which emphasize the importance of differentiating vesicular from non-vesicular components and establishing cargo localization ^4^, we characterized the protease-accessible proteome of sEV preparations from *Salmonella*-infected macrophages. Proteinase K effectively depleted accessible host proteins while largely preserving luminal EV markers and bacterial antigens. Unexpectedly, proteolysis enhanced rather than diminished the inflammatory activity of the preparations, suggesting that externally accessible proteins can influence how infection-associated vesicles stimulate recipient cells.

Protease accessibility does not establish whether a protein is integrated into the vesicle membrane, loosely associated with its exterior, part of a protein corona, or present in co-isolated material. Previous studies have used protease protection assays to distinguish surface-associated from encapsulated EV cytokines ^33^. By combining quantitative proteomics, peptide-level analysis, and normalization-independent assessments of protein retention, our study extends this approach to the broader infection-associated sEV proteome. Retention of TSG101 and ALIX in particle-normalized immunoblots supports preservation of protected vesicular components after proteolysis, independently of proteomic normalization.

TNF-α provides a particularly informative example. The sensitivity of its 26-kDa membrane-bound precursor is consistent with its type II transmembrane topology, which exposes the C-terminal extracellular domain, and with previous reports of membrane-associated TNF-α on sEVs ^34^. The 17-kDa soluble form was also sensitive to digestion, although its association with the vesicle fraction remains uncertain and could reflect receptor binding ^35^, adsorption, or co-isolation. Notably, retention of a cytoplasmic TNF-α peptide despite loss of immunoblot signal is consistent with cleavage of the extracellular antibody epitope while a protected intracellular fragment persists.

Protease sensitivity was also observed for CSF2RA, CD40 and MIF, and IL-1β and IL1RAP were detected in all three untreated replicates but not after proteinase K treatment. The detection of IL-1β in RAW264.7-derived sEVs is striking, since these macrophages lack the inflammasome adaptor ASC and cannot carry out canonical inflammasome-dependent maturation of IL-1β. They can, however, release unprocessed pro-IL-1β by a caspase-1-independent route ^36^, although whether that pathway operates during *Salmonella* infection has not been established. Our data raise a new possibility that association with extracellular vesicles provides an additional route by which the precursor leaves the cell, independently of inflammasome activation, a route that would be notable given that pro-IL-1β lacks a signal peptide and cannot use the conventional secretory pathway that some other proteins do. Consistent with this difference in inflammasome competence, sEVs from *Salmonella*-infected macrophages induced IL-1β release from recipient human THP-1 macrophages but not from RAW264.7 cells in our earlier work ^37^. Both identified IL-1β peptides (residues 134–144 and 192–205; **Table S3**) lie in a region shared by the precursor and the mature cytokine, so the proteomic data cannot discriminate between them, although the ASC deficiency of this line makes the precursor the more likely species. Loss of detectable IL-1β following digestion is consistent with protease accessibility, although its precise association with the vesicle preparation remains still unresolved. In contrast, CXCL10, CCL2, CCL9 and CCL5 showed no significant depletion after Proteinase K digestion, suggesting that protease susceptibility varies among immune mediators and some cytokines may be inside the lumen of the EV while others are externally exposed.

Cytokine localization on sEVs may also depend on the model used and source of the vesicles. A survey of 33 cytokines across eight human systems reported TNF-α to be predominantly encapsulated in most matrices but surface-associated in plasma ^33^, and found cytokine topology to vary with biological context and activation state. Moreover, one study identified IL-3 to be carried on the surface of EVs produced by Th2 cells, confirmed by proteinase K digestion ^8^.

Direct comparison is limited, since polymer-based precipitation used in that study ^33^, size exclusion chromatography separation ^8^, and ultracentrifugation (used here) recover different vesicle populations and different amounts of co-isolated protein, and the studies differ in species, cell model and protease. These findings caution against assigning a universal vesicular localization to individual cytokines.

Two recent studies place specific proteins on the surface of infected-macrophage EVs by indirect means, and our data test those assignments directly. First, a proteomic study of EVs from *S.* Typhimurium-infected J774A.1 macrophages identified MMP-9 as the most strongly induced matrisome protein and inferred its membrane localization from co-immunoprecipitation with LRP-1, fibronectin and annexin A2, alongside enrichment of CD14 ^28^. Second, a study of *Salmonella* sepsis used immunoelectron microscopy to place TfR, CD91 (LRP-1) and CD163 on the vesicle surface, where they capture circulating iron-containing proteins and restrict bacterial iron acquisition, with enrichment actively directed by Rab31 downstream of ER stress rather than passively inherited ^21^. In our preparations, LRP-1 (log₂FC −1.32, 62 peptides), fibronectin (−1.76), TfR (−1.25, 27 peptides) and CD14 (−3.35) were all removed by external protease, while annexin A2 was unchanged and CD163 was not detected, which could be due to sensitivity issues. MMP-9 was absent from our data, but MMP-14, a membrane-anchored metalloproteinase, was lost after digestion and carries topological evidence of surface exposure. Despite different isolation schemes, *Salmonella* strains and macrophage sources, LRP-1 has now been placed on the infected-macrophage vesicle exterior by co-immunoprecipitation, immunoelectron microscopy and protease shaving. The same enrichment of matrix-remodeling proteases appears in EVs from *Yersinia pestis*-infected human neutrophils ^20^, indicating that this is not specific to *Salmonella*.

Most detected *Salmonella* proteins resisted proteinase K digestion, and several were significantly increased relative to the surviving proteome. These increases are most plausibly compositional, since equal protein mass was analyzed after removal of a large fraction of accessible host protein; they indicate a greater share of what remains rather than more bacterial protein per vesicle. The persistence of most bacterial proteins after extracellular proteolysis indicates that infection-derived sEV preparations retain potentially antigenic material. Consistent with this, such vesicles carry bacterial antigens, induce pathogen-specific adaptive immunity⁶ and protect mice against lethal challenge ^22^, with similar immunogenic properties reported for sEVs from *Mycobacterium tuberculosis*-infected cells ^38^. Our findings extend these observations by demonstrating the proteolytic protection of *Salmonella*-derived proteins in infection-associated sEV preparations.

Unexpectedly, removal of surface-accessible proteins enhanced the immunostimulatory activity of sEV preparations despite depletion of several host inflammatory mediators. The post-digestion flow-through was inactive, indicating that the enhanced response remained with the vesicle fraction rather than released peptides. TNF-α neutralization reduced the enhanced *Il6* response to untreated levels, consistent with TNF-α-dependent amplification despite depletion of vesicle-associated TNF-α, and polymyxin B abolished the enhancement, implicating polymyxin-sensitive bacterial components. Bacterial LPS has been identified within infection-derived EVs and implicated in macrophage inflammatory signaling ^37,39^. In particular, Hui et al. demonstrated that *Salmonella*-associated exosomes induce TLR4-dependent TNF-α production, which was only partially inhibited by polymyxin B^37^. Although those experiments support intravesicular LPS, our findings do not establish whether proteolysis alters LPS exposure, vesicle uptake, or downstream signaling. Because proteinase K does not degrade LPS, the increased inflammatory response may result from changes in ligand accessibility or presentation following removal of associated proteins. The combined findings are therefore consistent with a model in which surface-accessible proteins influence recognition of pre-existing bacterial inflammatory ligands, with subsequent signaling amplified through TNF-α-dependent pathways. A pathogen can also limit EV inflammatory capacity directly, as *Y. pestis* does by using type III secretion effectors to exclude antimicrobial and proinflammatory cargo from neutrophil EVs ^20^. Our data show a different arrangement as the inflammatory potential is present within the vesicle but constrained by the vesicle’s own surface proteins. In one case the pathogen prevents the signal from being packaged; in the other the signal is packaged but not readily accessible.

This interpretation relates to emerging evidence that the EV protein corona influences vesicle interactions with recipient cells. One study reported enhanced uptake of corona-coated EVs by immune cells ^40^, and another study observed no substantial effect of such corona on association with phagocytic cells ^9^. Both studies examined EV coronas acquired from extracellular fluids, whereas we removed proteins already present on vesicles generated during infection. Proteinase K also depleted ITGAV (−1.90) and ITGA5 (−1.36), and because EV-surface integrins direct interactions with recipient cells ^41^., their removal could alter vesicle docking or uptake independently of ligand accessibility. The enhanced response may therefore reflect changes in binding, uptake or trafficking rather than a single mechanism.

Several limitations should be considered. Protease accessibility alone cannot distinguish membrane topology from co-isolation, and surface labeling, detergent-assisted digestion and density-gradient fractionation will be needed to refine cargo localization along with other methods. In the functional experiments, vesicle uptake was not directly measured and the two inhibitors were not tested in combination. Extending these observations in vivo will also be necessary to establish whether surface accessibility shapes vesicle activity in an intact host. Together these limitations prevent definitive assignment of the enhanced inflammatory response to a particular ligand or mechanism.

Collectively, our findings reveal distinct patterns of proteolytic accessibility among host and bacterial proteins associated with sEV preparations from *Salmonella*-infected macrophages. While many host membrane proteins and inflammatory mediators were accessible to extracellular proteolysis, bacterial proteins were predominantly protected. Unexpectedly, depletion of surface-accessible proteins enhanced inflammatory signaling, implicating the extracellular protein environment as a regulator of infection-associated vesicle immunostimulation. These findings highlight protein accessibility, rather than cargo abundance alone, as an important determinant of how infection-derived vesicle preparations interact with recipient immune cells.

## Methods

### Bacterial and mammalian cell culture

*Salmonella enterica* serovar Typhimurium (strain UK-1) was grown overnight at 37 °C with orbital shaking at 250 rpm in LB Miller medium. Mid-log cultures were prepared to an optical density of approximately 0.50 at 600 nm. Bacteria were pelleted at 6,000 x g for 10 minutes and washed twice in sterile 1× Phosphate-buffered saline (PBS). Bacterial concentration was adjusted to inoculate cells at a multiplicity of infection (MOI) of 5:1.

RAW264.7 murine macrophages (ATCC TIB-71) were cultured in Dulbecco’s Modified Eagle Medium (DMEM; Gibco 11-995-065) supplemented with 10% fetal bovine serum and 1% penicillin/streptomycin at 37 °C and 5% CO₂. On the day of infection, medium was replaced with DMEM containing 1% EV-depleted FBS (heat-inactivated FBS ultracentrifuged at 100,000 × g for 18 h) prior to the gentamicin protection assay ^6,22,26^. Here, bacterial suspensions, or an equal volume of sterile 1× PBS for mock conditions, were co-incubated with macrophages at MOI 5:1 for 2 hours at 37 °C and 5% CO₂. Media were removed and cells treated with 100 µg/mL gentamicin sulfate (30-005-CR) for 1 hour, after which DMEM containing 20 µg/mL gentamicin was applied for 24 hours.

### sEV isolation and analysis

The sEVs were isolated as we previously described^6,42,43^. Conditioned media were treated with protease inhibitor cocktail (PIA32953) and centrifuged for 10 minutes at 500 × g, 10 minutes at 4,000 × g and 30 minutes at 16,000 × g. The supernatant was passed through a 0.22 µm polyethersulfone filter. sEVs were isolated by ultracentrifugation at 100,000 × g for 180 minutes, performed twice, using an SW 32 Ti swinging-bucket rotor. The supernatant was discarded and the pellet resuspended in 1× protease inhibitor cocktail. sEV size and concentration were determined on a ZetaView QUATT (Particle Metrix) at 85 shutter and 100 gain. Samples were diluted in HPLC-grade water and injected to obtain median size and original concentration.

### sEV disruption

Three independent approaches were compared. For mechanical disruption, preparations were probe-sonicated with an ultrasonic homogenizer for 30 s, repeated twice, and maintained on ice between cycles. For thermal disruption, preparations were incubated at 95°C for 10 min and cooled immediately on ice. For enzymatic shaving, sEVs were incubated with proteinase K (Thermo Fisher 17916) at final concentration of 20 µg/mL in PBS for 1 h at 37 °C with gentle mixing every 15 min. Protease activity was quenched with phenylmethylsulfonyl fluoride (PMSF) at a final concentration of 100 µM followed by a further 15 min incubation. Samples were then concentrated on Amicon Ultra centrifugal filter units (MilliporeSigma) to remove proteinase K, low-molecular-weight digestion products and residual peptides. Untreated controls were processed identically, including PMSF addition and concentration, but without proteinase K.

### MACSPlex multiplex bead assay

Surface-marker profiling used the MACSPlex EV Kit IO, mouse (Miltenyi Biotec 130-122-211), which resolves 37 surface epitopes plus two isotype control bead populations, with the combined CD9/CD63/CD81 APC detection reagent. Vesicle input was standardized by NTA to 1 × 10¹⁰ particles/mL in 120 µL of kit buffer. Assays followed the manufacturer’s overnight protocol in 1.5 mL tubes. Briefly, 15 µL of capture beads were incubated with each sample overnight on an orbital shaker at 450 rpm. Beads were washed with 500 µL buffer (5 min, 3,000 × g), incubated for 1 h with 15 µL of APC-conjugated anti-CD9/CD63/CD81 detection reagent with shaking, washed again, and transferred to a 96-well round-bottom plate. Each run included a buffer-only blank containing capture beads, processed identically.

Blank signal was subtracted from the corresponding bead in each sample of the same run. A bead population was scored as detected when its raw signal reached at least twice its own blank, a bead-specific criterion used because background varied by more than an order of magnitude between populations; both isotype controls fell below this limit in every sample. Corrected values below the limit were reported as not detected rather than as negative values.

Treatment effects were expressed for each marker as the log₂ ratio of treated to matched untreated sample from the same preparation, which internally controls for vesicle input. Ratios were computed only where the untreated sample exceeded the detection limit; where a treated sample fell below it, the limit was substituted for the numerator and the result reported as an upper bound, so reported mean log₂ fold changes are geometric means. Treatments were then compared with untreated controls by two-tailed one-sample *t* tests of the per-replicate log₂ ratios against zero. The disruption-method comparison (**Fig. S1A**) used four preparations, two infected and two uninfected, with infection status in the model so each was compared against its own untreated control. The proteinase K experiment (**Fig. 1C**) used two infected preparations, one digested twice, giving three digestions in total. *P* values were adjusted across markers by Benjamini–Hochberg and are reported as *q* values, with significance at *q* < 0.05.

### Western blotting

Treated and untreated sEVs were concentrated on Amicon 10 kDa molecular weight cut-off filter units (MilliporeSigma). Protein from sEVs, cell pellets and conditioned medium was prepared by methanol–chloroform precipitation and quantified by bicinchoninic acid (BCA) assay (BioRad) and equal amounts of protein were loaded per lane for input material. However, for sEV samples, loading was normalized by particle number, with 4.5 × 10⁹ particles loaded per lane. Samples were reduced in 2-mercaptoethanol and separated on 4–12% bis-tris gels (Bio-Rad) at 75 V for 15–30 min followed by 175 V for 45 min, then transferred to PVDF membrane. Membranes were blocked in 1% milk and probed overnight at 4°C with anti-TNF-α (Novus NB600-587), anti-CD9 (System Biosciences), anti-TSG101 (System Biosciences), anti-ALIX (Santa Cruz) or anti-*Salmonella* O/H antigen (Thermo Fisher PA1-20811). HRP-conjugated goat anti-rabbit secondary antibody (Thermo Fisher 31460) was applied at 1:500 for 1–2 h with shaking. Clarity Max Western ECL substrate was applied for 5 min before imaging. PVDF membranes were not cut prior to antibody incubation. Western blot images were acquired digitally using a Bio-Rad GelDoc imaging system. The original, uncropped images are provided in the Supplementary Information (**Fig. S5-S6**).

### Cell culture-based functional studies

Naïve RAW264.7 macrophages were seeded in 6-well plates 24 h before stimulation. Medium was replaced with complete medium containing 2 µg/mL anti-IgG isotype control (BioLegend 400431), 2 µg/mL anti-TNF-α neutralizing antibody (BioLegend 506331), or 6.60 µg/mL polymyxin B sulfate (Thermo Fisher), and pre-incubated for 2 h at 37 °C and 5% CO₂. Cells were then co-incubated for 4 h with 1 µg of untreated sEVs, proteinase K-treated sEVs, proteinase K flow-through, or vehicle, before harvesting for RNA extraction. For reporter assays, QUANTI-Blue NF-κB/AP-1 reporter RAW264.7 cells (InvivoGen) were co-incubated for 24 h before supernatant collection and SEAP detection per manufacturer’s instructions.

### Gene expression analysis

Total RNA was extracted using the RNeasy Plus Micro Kit (Qiagen) according to the manufacturer’s instructions. Purified RNA was eluted in a final volume of 20 µL, and RNA concentration was determined using a NanoDrop UV–Vis spectrophotometer. Subsequently, 1 µg of total RNA was reverse-transcribed into complementary DNA (cDNA) using MultiScribe Reverse Transcriptase (Applied Biosystems). Quantitative real-time PCR (qRT-PCR) was performed using SYBR Green qPCR Master Mix (Bio-Rad Laboratories) to determine the relative mRNA expression levels of *Il6*. *β-Actin (Actb)* was used as the endogenous reference gene for normalization. The forward and reverse primer sequences (5′–3′) were as follows: *Actb*, GATGTATGAAGGCTTTGGTC and TGTGCACTTTTATTGGTCTC; and *Il6*, AAGAAATGATGGATGCTACC and GAGTTTCTGTATCTCTCTGAAG, respectively. Relative gene expression was calculated using the comparative quantification cycle (Cq) method. Cq values of target genes were normalized to those of *Actb*, and fold changes relative to the control group were determined using the 2^−ΔΔCq method.

### Mass spectrometry analysis

sEVs were analyzed at 10 µg per sample, quantified by microBCA assay (Cat#23235, Thermo FisherScientific). For digested samples, Amicon Ultra centrifugal filter units (50 kDa molecular weight cut-off) were used to remove proteinase K and digestion products. Three biological replicates per condition were separated by SDS-PAGE on 4–12% bis-tris gels (Bio-Rad). Lanes were excised, diced into 1 mm² cubes, and subjected to in-gel trypsin digestion as previously described^6,44^. Trypsin-digested peptides (200 ng) were loaded onto a 50 cm C18 capillary column (Thermo PepMap ES75500PN) on a Thermo Vanquish Neo UHPLC system. Peptides were separated over a 60 min gradient of solvent B (80% acetonitrile, 0.1% formic acid): 4–5% over 0.5 min, 5–28% over 44 min, 28–42% over 4 min, 42–55% over 4 min, then a ramp to 99% over 0.5 min and a 7 min hold for column washing. Eluted peptides were analyzed on a Thermo Orbitrap Exploris 240. MS1 scans were acquired at 120,000 resolution over 375–1,475 *m/z* with an AGC target of 3 × 10⁶. MS2 spectra were acquired using a TopN method within a 3 s cycle with HCD at 30% normalized collision energy, 15,000 resolution, an AGC target of 5 × 10⁵ and dynamic exclusion of 45 s. Raw LC-MS/MS data were processed in Proteome Discoverer 3.2.0.450 (Thermo Fisher). Spectra were searched with Sequest HT and Comet against UniProt reference proteomes for mouse (UP000000589) and *Salmonella enterica* serovar Typhimurium LT2 (UP000001014), along with a common contaminants database. Searches used full trypsin specificity with up to three missed cleavages. For Sequest HT, precursor and fragment mass tolerances were 10 ppm and 0.02 Da. Carbamidomethylation of cysteine (+57.021 Da) was set as a static modification; methionine oxidation (+15.995 Da) and protein N-terminal acetylation (+42.011 Da) were variable. Peptide-spectrum matches were validated with Percolator using a concatenated target–decoy strategy, with strict and relaxed FDR thresholds of 1% and 5% for peptide-spectrum matches, peptides and proteins. Only high-confidence peptides and master proteins were retained, and protein grouping used strict parsimony. Label-free quantification used Minora feature detection and the Precursor Ions Quantifier, based on precursor intensity from unique and razor peptides with a minimum replicate-feature requirement of 50%. Trypsin protein, introduced during in-gel digestion, was removed prior to statistical testing.

Peptide-level quantitation was exported from Proteome Discoverer as a PeptideGroups table and processed in R 4.4.1 using the QFeatures framework, following the Bioconductor expression-proteomics workflow of Hutchings et al.^45^ with the modifications described below. Because a retention-time alignment algorithm (match between runs) was applied during the identification search, processing began at the peptide level rather than the PSM level.

Of 12,527 peptide groups, entries were removed if they lacked a master protein accession (n = 35), matched a protein group containing a contaminant (n = 1,136), were flagged in the Quan Info field as No Quan Values, Shared or Not Reliable (n = 1,601), mapped to more than one protein group, or were not of high confidence, leaving 9,755 peptide groups.

Missing values were strongly condition-dependent, comprising 3.9–8.9% of control measurements and 32.1–39.3% of proteinase K measurements, and increased monotonically with decreasing intensity. As proteinase K removes surface-exposed protein by design, a global missing-value filter would have discarded precisely the peptides carrying the treatment effect. Peptides were therefore retained if quantified in at least two of three replicates of either condition (9,732 peptides) and classified by the type of missingness. Peptides quantified in at least two of three replicates of both conditions (n = 6,564) were treated as missing at random and imputed by k-nearest neighbours; peptides absent from an entire condition (n = 3,168) were treated as missing not at random and imputed with a per-sample detection floor set at the first percentile of observed intensities in that sample. The floor was derived from the full matrix rather than the censored subset, since the observed values of these peptides derive only from the condition in which they were detected. Imputation was performed on log2-transformed data and replaced 11,925 missing values.

Peptides were aggregated to master protein accessions by robust summarization (MsCoreUtils)^46^. Protein-level abundances were normalized by median centering, and differential abundance between conditions was assessed with limma^47^ using an empirical Bayes moderated t-test with an intensity-dependent prior variance trend and robust hyperparameter estimation. P values were adjusted by the Benjamini–Hochberg method. Proteins supported by fewer than two peptides were excluded from reporting. Proteins detected in no control replicate (n = 3) were excluded as imputation artefacts.

#### Assessment of normalization and vesicle integrity

Because proteinase K removes a substantial fraction of total protein, median centering may introduce compositional bias by assuming that most proteins remain unchanged. We hence applied two normalization-independent measures to assess the preservation of protected vesicular components. First, we calculated within-sample peptide abundance percentiles, which are invariant to sample-level scaling. Second, peptide retention was defined as the proportion of peptides detected in at least two untreated replicates that remained detectable in at least two treated replicates, independently of intensity measurements.

Normalization sensitivity was evaluated using eight schemes applied to the same un-normalized protein matrix, so that the offsets are directly comparable: no normalization, median centering (primary), median of ratios, trimmed mean of M values, total signal, a data-driven stable-rank reference set (n = 519), 20 candidate luminal EV reference proteins, and 11 *Salmonella* outer membrane and envelope proteins. The stable-rank set comprised the proteins with the lowest variability in within-sample abundance rank across the six samples. The luminal set comprised proteins associated with endosomal sorting, vesicle trafficking and EV-associated compartments (**Table S5**). The *Salmonella* reference comprised every bacterial protein in the dataset except flagellin. Each scheme defines a six-element vector of per-sample shifts, centered to mean zero, which was subtracted before refitting *limma*; proteins were then classified using the primary reporting rules. No normalization and the *Salmonella* reference are reported as bounds on the absolute scale rather than as candidate strategies. Normalization was applied after imputation, so all schemes share the same imputed values.

#### Annotation of surface exposure

Topological annotations were retrieved programmatically from the UniProt REST API for every master protein accession in the dataset, using the transmembrane, topological domain, lipidation and subcellular location fields. A protein was classified as carrying topological evidence of surface exposure if it has an annotated GPI anchor, or a transmembrane segment together with an annotated extracellular topological domain. Proteins with curated plasma membrane localization but no such topology were not used, because that annotation is assigned whenever cell membrane appears anywhere among multiple subcellular locations and in this dataset captured proteins with no plausible route to the vesicle exterior.

#### Peptide-level analysis of TNF-α

Peptides mapping to mouse pro-TNF-α (UniProt P06804, 235 aa) were assigned to topological domains using the annotation for that entry: cytoplasmic, residues 1–35; transmembrane, 36– 56; extracellular, 57–235. To remove any dependence on sample loading or normalization, the abundance of each extracellular-domain peptide was expressed as a log2 ratio to the cytoplasmic-domain peptide measured in the same injection.

## Statistics and software

Figures were prepared in GraphPad Prism 10 (GraphPad Software) and in R using *ggplot2*. For functional assays, group differences were assessed by one-way ANOVA with Tukey’s post hoc test; groups not sharing a letter differ significantly at *P* < 0.05. Significance is denoted n.s. *P* > 0.05, \**P* < 0.05, \*\**P* < 0.01, \*\*\**P* < 0.001, \*\*\*\**P* < 0.0001. Illustrations and assay schematics were generated with BioRender (BioRender.com).

## Data availability

Images were acquired digitally using a Bio-Rad GelDoc imaging system. The mass spectrometry proteomics data have been deposited to the ProteomeXchange Consortium via the PRIDE partner repository with the dataset identifier PXD085270. Reviewer token: VMXaZ1I5zdVa.

## Ethics Statement

This study did not involve human participants or experiments on live animals. Institutional human or animal research ethics approval was not required.

## Author Contributions

JJC – Design of experimental approaches, execution of assays, data and statistical analyses, interpretation of results, manuscript writing, figure design

CL – Execution of assays, figure design, manuscript writing

SE – Execution of assays, manuscript writing

XW – Experimental design, mass spectrometry acquisition and analysis, manuscript writing

MJF – Conceptualization and study design, data analysis, figure preparation, interpretation of results, manuscript review and editing, supervision, funding acquisition

## Acknowledgements

This work was supported by the National Institute of Allergy and Infectious Diseases of the National Institutes of Health under award number R01AI158749 (to M.J.F.).

